# Notch2 signaling instructs viral and bacterial TLR responsiveness in B cells

**DOI:** 10.1101/2025.10.10.681480

**Authors:** Jennifer Londregan, Isaiah Rozich, Brian T. Gaudette, Nicholas J. Tursi, David B. Weiner, Ivan Maillard, David Allman

## Abstract

Marginal zone (MZ) B cells are hyperresponsive to bacterial Toll-like Receptor (TLR) ligands. However, the full extent of TLR responsiveness for MZ B cells and the mechanisms regulating such responses are unclear. We report that Notch2 activation establishes MZ B cell responsiveness for the viral dsRNA receptor TLR3 and augments responses for the LPS receptor TLR4. Notch2 ligation accelerated Myc induction, mitosis, and plasma cell differentiation to LPS. Further, TLR3 expression in MZ B cells was Notch2 dependent, and ectopic Notch2 signaling was sufficient to promote robust TLR3 responsiveness dependent on the TIR-domain-containing adapter TRIF and the kinase BTK. TLR3 engagement in MZ B cells promoted proliferation, differentiation, and the secretion of IgG2b and IgG2c antibodies. Our results establish a novel role for Notch2 in establishing TLR3 and TLR4 responsiveness in B cells and suggest that MZ B cells play unappreciated roles in immunity against RNA viruses.

## Introduction

Leukocytes use Toll-like receptors (TLRs) to sense and respond to an array of conserved bacterial and viral ligands. While macrophages and other innate cells rely heavily on pattern recognition receptors, lymphocytes also employ TLRs to initiate or augment responses to microbial antigens^1,2,3,4,5,6,7^ Among B cells, both marginal zone (MZ) and “B1” B cells are especially responsive to TLR ligands such as lipopolysaccharide (LPS), the main ligand for TLR4^8^. MZ B cells are naïve B cells that reside in and around the MZ of the spleen where they are uniquely poised to generate large numbers of antibody-secreting plasma cells in response to blood-borne bacterial antigens^9,10,11^. Many of the unique properties of MZ B cells are controlled by the Notch2 transmembrane receptor^12,13^.

Little is known about the earliest phases of B cell responses and the genetic, epigenetic, and biosynthetic processes governing cell cycle entry and plasma cell differentiation.One key outcome of lymphocyte activation is multiple rounds of mitosis, which serves to grow the size of responding cell populations and often permits the cell-intrinsic transition from one cell state to another, such as from naïve lymphocyte to effector. For instance, most B cells must divide several times before yielding plasma cells^14^. MZ B cells on the other hand are able to generate plasma cells despite full blockade of mitosis^12,15^.

Development and maintenance of the MZ B cell pool is uniquely and strictly dependent on continuous ligation of the Notch2 receptor by the Delta-like-1 (DL1) ligand on splenic fibroblasts^16,17,18,13^. DL1-Notch2 interactions result in two sequential proteolytic cleavage events of the Notch2 receptor that release its intracellular domain to translocate to the nucleus where it regulates the transcription of large numbers of genes^19,20,21^. Recent data indicate that MZ B cells use Notch2 to regulate two functionally distinct collections of genes, Myc-dependent target genes enriched for cell growth regulators, and Myc-independent targets that the positioning of B cells in and around the MZ^13^. However, the relevance of Notch2 signaling to other unique functional properties of MZ B cells is unclear.

TLR3 is a primarily endosomal TLR activated by dsRNA of at least 40 base pairs^22^, a common intermediate produced during the replication of RNA viruses^23^ that can be mimicked with the synthetic analog poly(I:C)^24,25^. Whereas most TLRs require the adapter protein MYD88, TLR3 and TLR4 are regulated by the alternative adapter TRIF; TLR3 uses TRIF exclusively whereas TLR4 uses both TRIF and Myd88^26,27^. Additionally, phosphorylation of the TIR domain of TLR3 critically depends on Bruton’s tyrosine kinase (BTK)^28,29^,which functions downstream of the B cell receptor and is essential for optimal B cell clonal selection and survival^30,31^. TLR3 function is important for a variety of innate cells but has no obvious direct role for T or B cells. Notably though, MZ B cells are unique among B cells in that they possess *Tlr3* transcripts^32,5^.

We demonstrate here that Notch2 activity is critically tied to TLR4 and TLR3 responsiveness in B cells, while also accelerating cell cycle entry and the overall proliferative burst in response to ligation of either receptor. Specifically, DL1 stimulation of Notch2-inexperienced follicular B cells significantly augmented responsiveness to poly(I:C) and low doses of LPS while also accelerating cell cycle entry and amplifying numbers of induced plasma cells in a manner mimicking freshly isolated MZ B cells. Additionally, *Tlr3* mRNA abundance was reduced for MZ B cells upon *in situ* Notch2 blockade, and DL1 stimulation augmented class switching and the secretion of IgG_1_, IgG_2b_, and IgG_2c_ antibodies of follicular B cells. Our results therefore reveal that Notch2 regulates TLR3 and TLR4 response programs in B cells and reveal the possibility that MZ B cells play poorly understood roles in protective responses to many RNA viruses.

## Materials and Methods

### Mice

B6.Blimp1^+/GFP^ (Prdm1^+/GFP^) mice obtained from Steven Nutt were bred and housed in our animal colony. cMyc GFP KI mice, C57BL/6 mice, CBA/carter and CBA.xid mice were obtained from Jackson Laboratories and were bred and housed in our animal colony at the University of Pennsylvania. Mb1cre mice and *Tlr3^f/f^* mice were obtained from Jackson Laboratories and crossed to generate Mb1cre *Tlr3^f/f^* animals. All animal procedures were approved by the University of Pennsylvania Office of Regulatory Affairs.

### Cell Culture

Spleens were harvested from mice, minced between frosted glass slides, and lysed with ACK lysis buffer to remove red blood cells. Single cell suspensions were labeled with Cell Trace Yellow or Cell Trace Violet, as indicated. B1 B cells were isolated from the peritoneal cavity of mice by peritoneal lavage with 10 mL of warm RPMI followed by agitation and gentle recovery of injected fluid. Follicular B cells were purified by CD23 positive selection using Miltenyi MACS columns. Marginal Zone B cells were purified by CD23 negative selection, followed by a CD21 positive selection of CD23-flow through using MACS columns. Purified cells were plated in 96-well plates at 2×10^5^ cells per well with culture media containing RPMI 1640 supplemented with 10% Fetal Bovine Serum, 1% penicillin streptomycin, L-glutamine, HEPES, non-essential amino acids, sodium pyruvate, and 0.1% 2-mercaptoethanol and gentamicin. Cells were stimulated with indicated concentrations of LPS (from salmonella enterica) or Poly(I:C), with or without 1 μM Ibrutinib for indicated time points ranging from 24-96 hours and incubated at 5% CO2 and 37 C. Other BCR signaling inhibitors such as R788 and R406 were used at 0.1 μM, and I406 was used at 1μM. Stimulation conditions were plated at culture initiation. OP9 and OP9-DL1 stromal cells (obtained from Warren Pear, University of Pennsylvania) were cultured and maintained in media containing alpha-MEM, 20% Fetal Bovine Serum, 1% L-glutamine, sodium bicarbonate, and 0.1% gentamicin. Cells were split at 60% confluency and plated out at 1.5×10^5^/well in monolayers 24 hours prior to B cell culture initiation.

### Flow Cytometry

Cells were removed from culture by gentle pipetting followed by treatment with FACS buffer containing 1% Bovine Serum Albumin and 0.5% 2mM EDTA. Cells were stained with surface antibodies and viability exclusion was performed with ToPRO-3. All flow cytometry was performed on a BD FACSSymphony A3 and analyzed with FlowJo v10 software. The data for this manuscript were generated in the Penn Cytomics and Cell Sorting Shared Resource Laboratory at the University of Pennsylvania (RRID:SCR_022376). Penn Cytomics is partially supported by the Abramson Cancer Center NCI Grant (P30 016520).

### Antibodies

CD138 (281-2)-BV-421[1:1000] was purchased from BioLegend. CD19 (1D3)-BUV661 [1:200] was purchased from BD Biosciences.

### Intracellular Staining

Intracellular pBTK staining were performed using BD Cytofix and BD Phosphlow Perm Buffer III two-step protocol. pBTK (N35-86)-PE[1:200] was purchased from Cell Signaling Technology.

### DNA/RNA Staining

DNA/RNA staining for cell cycle analysis was performed by first fixing cells to fresh chilled 70% ethanol dropwise while vortexing and incubating for 1 hour at 4 degrees Celsius. Cells were then washed and stained with DAPI at 1:1000 and incubated at room temperature for 30 minutes. Without washing, Pyronin Y was added at 20X for 10 minutes at room temperature, then run on the cytometer in linear. For analysis, doublet cells were rigorously excluded by an area/height doublet exclusion, followed by a DAPI area/DAPI height doublet exclusion.

### Calculation of Cell Numbers and Division Dynamics

To determine the number of cells in division and divorce the impact of cell expansion on division numbers, a calculation of mean division numbers were adapted from Heinzel S. et al and Hawkins ED et al^33,34^. FlowJo software was used to perform proliferation modeling by setting the undivided peak gate and estimating the number of division peaks to find the best fit. Values for the number of events in each division gate were then determined. Counting beads were used, gating on them by FSC/SSC to determine the number of events in the bead gate. With these values, the following procedure was followed to obtain mean division and division rate values.

Determine the <u>total cell number</u>: (# cell events/ # bead events) x (bead count/sample volume) x total volume in FACS tube
Determine the <u>fraction divided</u> by (# events in division n/ # total events) x total cell # computed from A (perform for each division gate).
Generate <u>precursor cohort values</u>: (# cells in division n/ 2^n.5^). Note that instead of using the division number (n) for the precursor cohort, assume on average each cell is half-way through any given division (i.e. for a division number of 2, use 2.5).
Determine the <u>total cohort #:</u> (C_1_+C_2_+C_3_…C_n_)
Estimate* the <u>mean division number</u> (*for when data are not normally distributed) ([C_1_ x 1] + [C_2_ x 2] + [C_3_ x 3]… [C_n_ x n])/ total cohort #
Graph mean division number values over time and determine <u>division destiny</u> by the maximum division number value. Determine <u>division rate</u> by performing a linear regression where the slope is equal to the division rate.

### ELISpot Assay

ELISpot assay plates (Millipore MSIPN4W50) were coated with capture goat anti Ig-heavy/Ig-light antibody (Southern Biotech 1010-01) at 10 μg/mL and blocked with complete growth media (RPMI, 10%FBS, HEPES, Penn/Strep, NEAA, Na-Pyruvate, 2-ME, L-glutamine). Cells were stimulated in tissue culture-treated plates with indicated stimulation conditions (LPS or Poly (I:C) at 10 mg/mL) for 72 hours at a cell concentration of 2×10^5^/well on OP9 or OP9-DL1 monolayers and then directly transferred to blocked ELISpot plates at the indicated serial dilutions and incubated overnight. Biotinylated goat anti IgM, IgG, IgG_1_, IgG_2b_, IgG_2c_, or IgG_3_ (Southern Biotech 1020-08, 1070-08, 1090-08, 1079-08, 1100-08) capture antibody was used at 0.1 μg/mL. Streptavidin alkaline phosphatase was used at 1:10,000 final dilution (Sigma Aldrich E2636). Spots were developed using BCIP/NBT liquid substrate and 1M sodium phosphate stop solution. Image capture was performed using the CTL Immunospot Analyzer and software (Cellular Technologies Limited). Counting was performed by eye and averaged by dilution for each condition in replicates of 3. Final images are displayed with sharpness level of 100%, brightness and contrast levels of 15% applied to every ELISpot image indiscriminately.

### RNA Sequencing

Cell populations were sorted directly into Trizol with 0.5% 2-ME and held at −80 °C until RNA preparation. RNA was prepared by published Trizol RNA-extraction protocol (Thermo-Fisher Scientific). RNA was co-precipitated using glycogen as a carrier. RNA was quantified using Qubit RNA high sensitivity fluorometric assay. cDNA was prepared using Takara Clonetech SMART-Seq® v4 Ultra® Low Input RNA Kit according to protocol using 500–1000 ng RNA as input. cDNA was quantified and qualified using HSDNA assay on an Agilent 2200 bioanalyzer. RNA-seq libraries were constructed using the Illumina Nextera XT kit with 150 ng cDNA input. Libraries were quality controlled and quantified by bioanalyzer and pooled at equal molar ratio preceding sequencing Illumina Hiseq (50 bp SE v4 high output) and Illumina Nextseq500 (75 bp SE v2) machines.

### Pseudoalignment and Gene Expression

Transcript abundance was computed by pseudoalignment with Kallisto^65^. Transcript per million (TPM) values were then normalized and fitted to a linear model by empirical Bayes method with the Voom and Limma R packages^66,67^ and differential gene expression was defined as a Benjemini and Hochberg corrected *p*-value of < 0.05 and fold change > 2 unless otherwise noted.

## Results

We first sought to establish a comprehensive comparison of LPS responsiveness in MZ versus follicular B cells. We began by evaluating Myc induction kinetics upon in vitro LPS stimulation using mice harboring a Myc-GFP fusion protein^35^. Compared to follicular B cells, basal Myc levels were higher in MZ B cells^12^. Upon stimulation, MZ B cells upregulated Myc significantly faster and maintained higher levels of Myc over 4h (**Figure 1A**). We performed a head-to-head comparison of LPS-induced proliferation kinetics in LPS-stimulated cells using purified Cell Trace Yellow (CTY)-labeled WT follicular and MZ B cells. Consistent with accelerated Myc up-regulation, a substantial fraction of MZ B cells experienced one or more rounds of mitosis within 40 hours, whereas by 65 hours the observed number of cell divisions was comparable for each (**Figure 1B**). We employed linear regression analysis to calculate mean division numbers as a function of time. Time to first division (μtd1, or [1-y-intercept]/slope) and overall time to complete subsequent divisions (1/slope) were significantly faster for MZ B cells (Δ17h between follicular and MZ B cells for μtd1, and a Δ10h for subsequent divisions) (**Figure 1C**).

**Figure 1.**
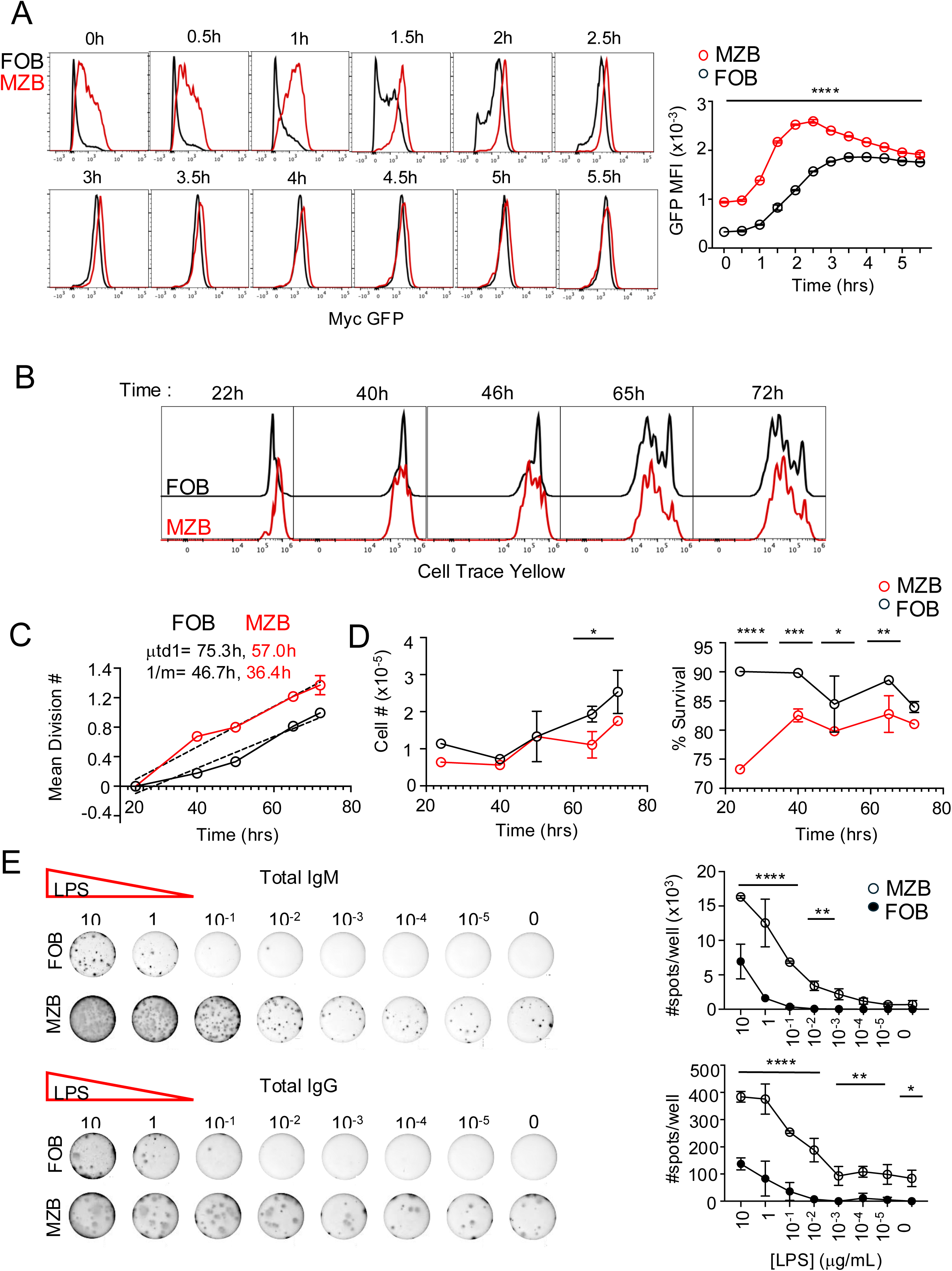
Marginal zone B cells’ LPS hypersensitivity owed to their more rapid response and plasma cell poising. Follicular (FO) or Marginal Zone (MZ) B cells were purified from Myc-GFP splenocytes by CD23 and subsequent CD21 selection and stimulated with 10μg/mL LPS and examined for Myc expression every half-hour for 5.5h. Myc-GFP histograms are displayed comparing MZ (red) and FO (black) B cells for Myc expression, MFI quantified to the right (A). Purified FO and MZ B cells were isolated, Cell Trace Yellow labeled and subsequently stimulated with 10μg/mL LPS over 22-72h. Histograms depict Cell Trace Yellow peaks between FO or MZ B cells (B). Mean division number and linear regression are depicted with values for time to first division (μtd1) and time to subsequent divisions (1/m) are displayed (C). Cell number over time in culture (left) and % of ToPRO-3^-^ cells over time in culture (right) are displayed (D). Total IgG and IgM ELISpots performed for purified FO or MZ B cells after 48h of stimulation in culture with indicated dose of LPS, 1:16 dilution is shown. Spots/well are quantified to the right (E). 2-way ANOVA performed, * = p<0.05, ** = p<0.005, *** = p<0.001 **** = p<0.0001. Triplicate cultures, repeated 3-5 times.

Quantification of cell numbers over time revealed a lower cell count of MZ B cells (left) despite enhanced division, which was consistent with overall lower survival at earlier time points (**Figure 1D**). Finally, as expected based on past results^11,12^ MZ B cells were predisposed to generate antibody-secreting plasma cells, including at low LPS doses (**Figure 1E**). Both IgM and total IgG secretion was observed with cells derived from the MZ pool, signifying their ability to undergo class-switch recombination (CSR). Altogether, these data demonstrate that MZ B cells respond robustly to LPS stimulation as individual cells undergo an earlier and more extensive proliferative and differentiative response than follicular B cells.

### Notch2 accelerates and amplifies LPS-driven proliferation

Given the unique dependency of MZ B cells on Notch2^18^, we asked whether Notch2 engagement was sufficient to enhance LPS responses. We co-cultured CD23^+^ CTY-labeled follicular B cells with OP9 stromal cells versus OP9 cells engineered to express the Notch ligand DL1^36^. Cultures were evaluated at multiple time points after LPS stimulation ranging from 26-93 hours, and absolute cell numbers quantified using counting beads added to samples immediately before analysis^34^. Notch2 ligation both accelerated the time with which LPS-stimulated B cells entered division and augmented overall burst size. As shown in **Figure 2A**, within 41 hours we observed evidence of cell division for DL1 + LPS stimulated cells, whereas only a small number of cells underwent division by 50 hours in control OP9 cultures. Moreover, overall viable cell recoveries were greater for OP9-DL1 co-cultures, apart from later time points when DL1-stimulated B cells experienced greater cell death, likely owing to depletion of nutrients or differentiation-induced death (**Figure 2B**).

**Figure 2.**
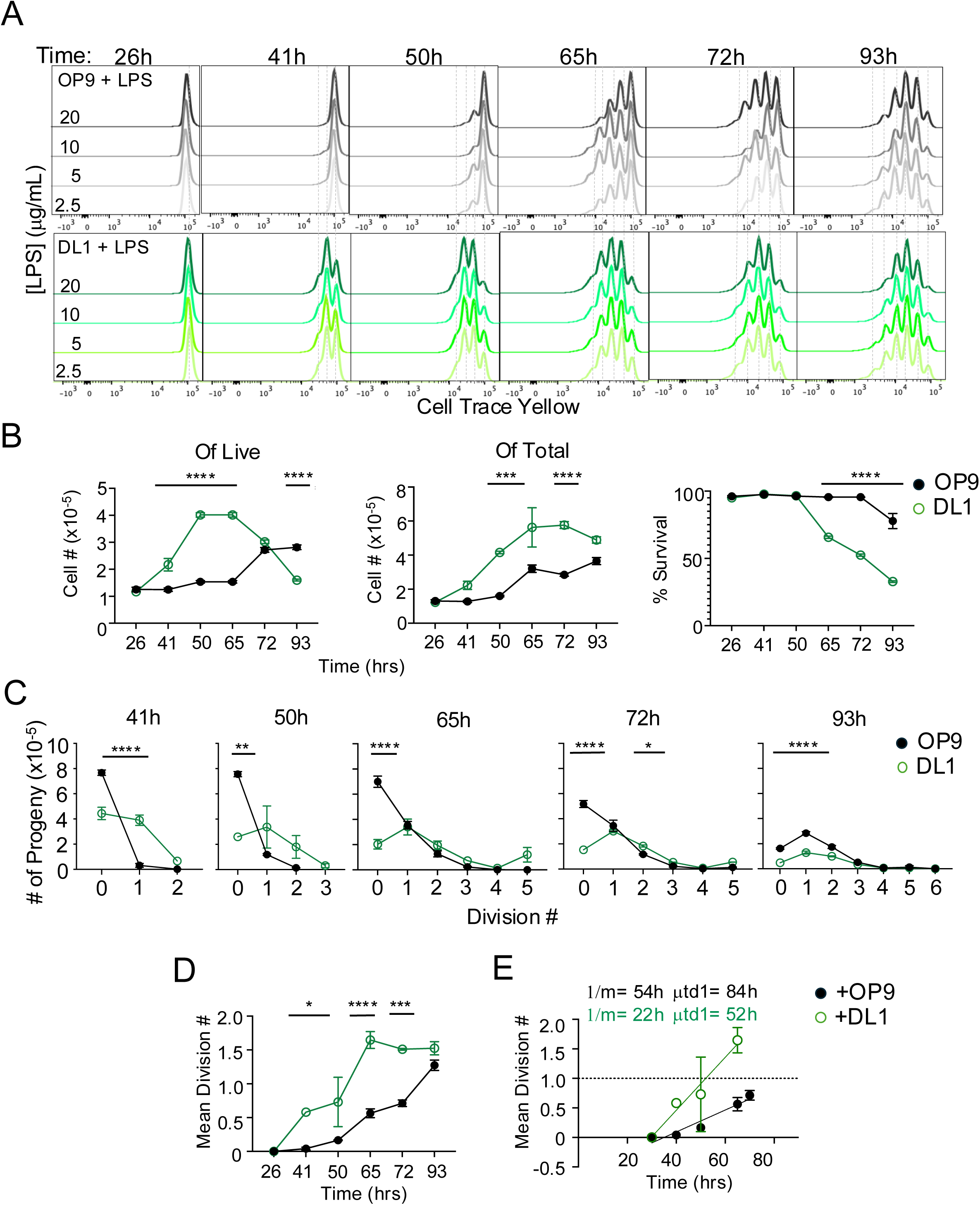
Notch2 augments the proliferative response to LPS for FO B cells through early Myc induction. Follicular B (FO B) cells, purified from splenocytes based on CD23 expression, were cell trace yellow labeled and co-cultured with OP9 stromal cells (grey) or OP9 stromal cells transduced to express Notch2-ligand DL1 (green). FO B cells were then stimulated with indicated amounts of LPS in culture for 26-93 hours, concentration denoted by increasingly darker shades of green or grey. Resulting histograms depicted (A). Cell number over time from live (left) and total (right) from cultures. All quantification shown at the 20 μg/mL dose (B). Percentage of ToPro-3 negative, viable cells over time in culture shown for OP9 (black) or DL1 (green). Precursor cohort plots depict the number of precursors in each respective cell division for OP9-experienced FOB cells (left, black) or DL1-experienced cells (right, green) at each dividing timepoint stimulated with 20 μg/mL LPS (C). The mean division number (MDN) over time of all B cells on OP9 (black) or DL1 (green) are depicted (D). Linear regression analysis was performed on the MDN over time to examine the average time to complete the first division (μtd1) and time to complete subsequent divisions, or 1/slope (E). 2-way ANOVA performed B-E, * = p<0.05, ** = p<0.005, *** = p<0.001 **** = p<0.0001. Results repeated in at least 5 independent experiments.

To assess the impact of DL1 stimulation on the progression of input cells across divisions more deeply, we used mathematical modeling to estimate the number of divisions experienced by each cell on average using a data analysis platform that separates the impact of cell division on the total number of cells in culture over time^33^ (see **Methods**). First, we generated precursor cohort plots to quantify the number of input cells that enter each round of division^37^. As shown in **Figure 2C**, at early time points a far greater fraction of B cells underwent one or more divisions when co-cultured on OP9-DL1 cells compared to OP9 controls. Next, we summed precursor cohorts for each sample to calculate mean division number (MDNs) for each time point, thus confirming that on average DL1-stimulated cells achieved a greater number of cell divisions (**Figure 2D**). We then performed linear regression analyses of MDN plots to obtain time to first division ([1-y intercept/]slope) and time between divisions (1/slope) for responding cells^33^. This revealed that LPS + DL1 stimulation decreased the mean time at which cells completed the first division (“μtd1”), and the time needed for responding cells to progress through each division (1/m), as these values were found to be 52h vs. 84h for the first division, and 22h vs. 54h for subsequent divisions. These results reveal that Notch2-experienced cells achieved on average a 32h ‘head start’ for cell division (**Figure 2E**).

To further test the notion that DL1-stimulated cells initiated cell division sooner, we examined the transition from the G_0_ to G_1_ phase of the cell cycle with and without DL1. We stimulated resting follicular B cells with 10 μg/ml LPS over 24-48h and measured the frequency of cells in each phase of cell cycle by staining with a DNA dye (DAPI) and a RNA dye (PyroninY).

Combining RNA and DNA content analyses allows resolution of cells in each phase of the cell cycle, including G_0_ (low RNA, 2N DNA) versus G_1_ (high RNA, 2N DNA)^38^. Cellular doublets were excluded from collection analyses using the width of the DAPI signal (see Materials and Methods). As shown in **Figure 3A**, at 24 hours a greater fraction of DL1-experienced B cells was detected in G_1_ compared to more B cells in G_0_ on OP9 control cultures. By 48 hours, B cells in OP9 cocultures had “caught up” to Notch2-experienced cells in G_1_-S, whereas DL1 cocultures exhibited a trending increase in the G_2_/M phase. Consistent with these data, Myc peaked earlier for LPS + DL1 stimulated cells at around 26h vs 50h without DL1 (**Figure 3B**).

**Figure 3.**
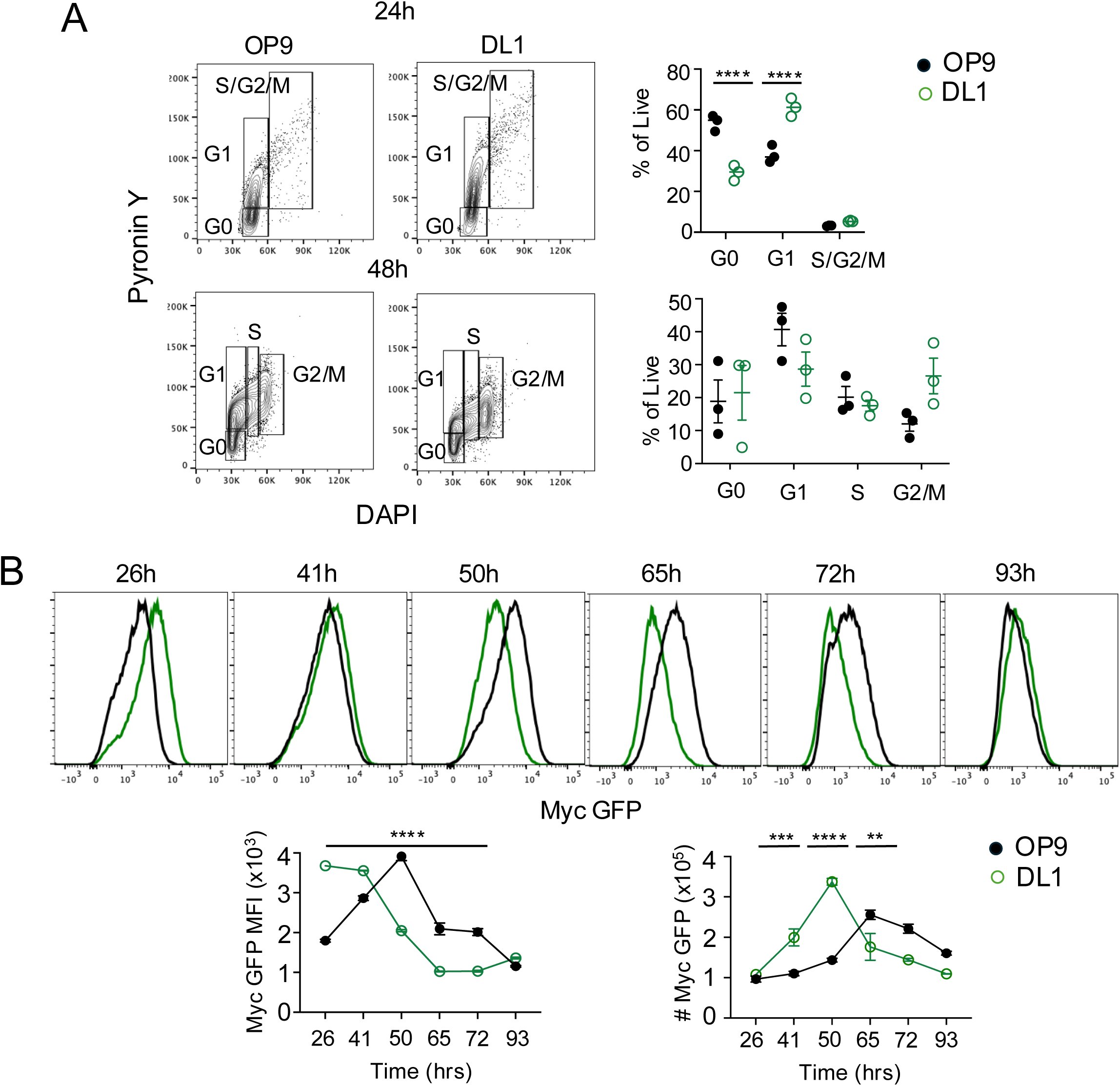
Notch2 speeds up division time and cell cycle entry for LPS-stimulated FO B cells. Purified FO B cells were co-cultured on OP9 or OP9-DL1 and stimulated with 10μg/mL LPS to examine DNA and RNA content and gauge cell cycle. Flow plots depict DAPI vs. Pyronin Y for cells in G0, G1, S, and G2/M phases at 24h or 48h (A). Purified FO B cells from Myc GFP reporters were co-cultured on OP9 or OP9-DL1 and stimulated with 20μg/mL LPS, Histograms of cMyc-GFP signal between OP9 (black) or DL1 (green) B cells stimulated with 20 μg/mL over indicated timepoints are shown. Quantification of cMyc GFP MFI over time between OP9 and DL1 (left) and the number of Myc+ cells over time (right) (B) Results are graphed on the right as a percentage of live single lymphocytes (B). 2-way ANOVA performed, * = p<0.05, ** = p<0.005, *** = p<0.001 **** = p<0.0001. Results repeated in 3-5 independent experiments.

We conclude that DL1 engagement drives LPS-stimulated B cells to accelerate their time to division and increase their proliferative capacity across several rounds of division.

### Notch2 accelerates pro-growth gene expression

To gain a mechanistic understanding of how Notch2 augments LPS-driven B cell responses, we used RNAseq to assess differentially expressed genes (DEGs) in viable follicular B cells from OP9 or OP9-DL1 co-cultures with 10μg/mL LPS stimulation for 20-, 44-, or 68-hours. We examined all DEGs with a log2 fold change greater than or equal to 1, or less than or equal to −1 at all three time points, combined each list and removed any duplicates. At 20 hours, we observed a substantial cluster of DEGs for DL1-stimulated cells with far fewer DEGs observed at later time points, signifying that many transcriptional changes are occurring early after Notch2 activation. Among the top DEGs at the earliest timepoint, multiple genes pertained to translation, metabolism, migration, proliferation, and viral pathways (**Figure 4A**). Gene set enrichment analysis similarly revealed early hallmark program changes in DL1-stimulated cells, including for canonical Notch targets and numerous pro-growth and division pathways such as Myc, E2F and G2M as well as PI-3-kinase/AKT/mTOR activity. Alternatively, pathways enriched in LPS-stimulated B cells without Notch2 at this time point included IFNγ, IFNα, TNFα, and other pathways classically associated with inflammation. At 44h, however, all those hallmarks were now enriched even without Notch2 signaling such as E2F, Myc, mTORC1, and the oxidative phosphorylation pathway, highlighting how DL1 accelerates this response (**Figure 4B**). Regarding specific genes, at 20 hours DL1+LPS stimulated cells exhibited reduced mRNA abundance for the plasma cell-repressive genes *Bach2* and *Bcl6* and increased transcript abundance for the pro-plasma cell differentiation genes *Prdm1* and *Xbp1* (**Figure 4C, column 1**). We examined several transcripts related to cell cycle, translation and ER stress, and antibody secretion as previously described in Gaudette et al 2021^12^ and found a modest upregulation *Cdk6*, *Top2a,* and *Aurka* transcripts at the early time point, without affecting B cell fate determining *Pax5*, or the ER chaperone gene *Hspa5*. Similarly, we verified that direct Notch2 targets like *Hes1* were upregulated in DL1-experienced B cells compared to controls. (**Figure 4C** **column 2, bottom**). We observed no significant change in expression to TLR-related genes such as the common adapter Myd88, or the LPS co-receptor Cd180, and Ticam1, the gene encoding the alternate TLR adapter TRIF. Most notably however, we found a robust early upregulation of *Tlr3* transcripts on DL1 after 20 hrs (**Figure 4B, column 2, top**). We conclude that Notch2 activity accelerates the induced transcription of a wide array of genes that promote cell growth and plasma cell differentiation, while also providing a unique potential mechanism for fostering TLR3 function in B cells.

**Figure 4.**
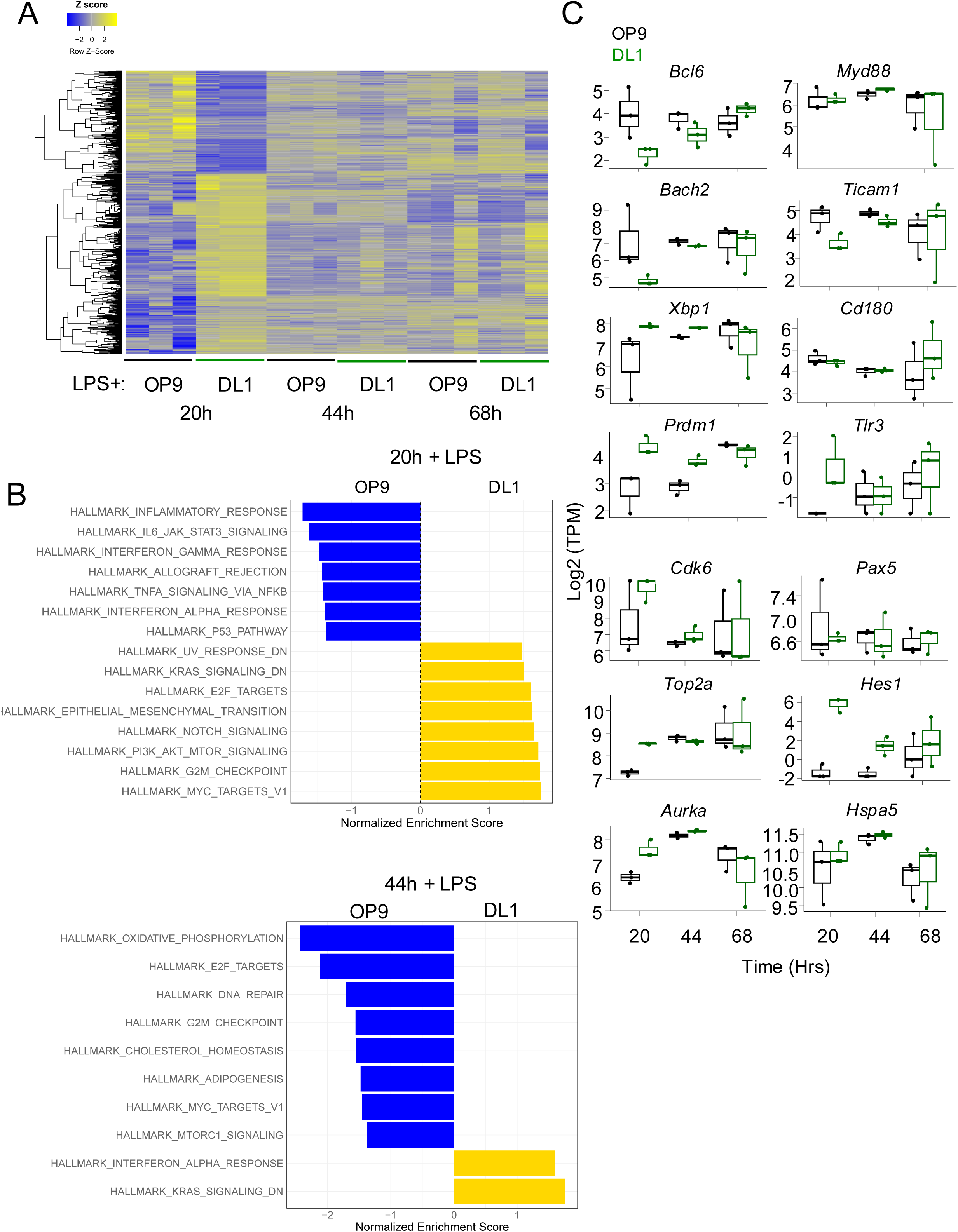
Forced Notch2 endows early transcriptional changes associated with activation, plasma cell differentiation, and TLR-responsiveness. Purified FO B cells were co-cultured with OP9 or OP9-DL1 for 20, 44, or 68h +/-10 μg/mL LPS and approximately 25,000 cells were subsequently sorted out of triplicate cultures for viability and purity directly into Trizol for bulk RNA sequencing. Shown is a heat map of differentially expressed genes over time, generated by all genes with a log2 fold change greater than or equal to 1 and less than or equal to −1 for each timepoint, then combining the 3 lists and excluding any duplicates, and normalizing gene expression per row (1482 genes total). Heat maps are expressed as a z score (A). Gene set enrichment analysis was performed using hallmark pathway analyses for indicated signatures comparing LPS stimulated FO B cells on OP9-DL1 to OP9 alone after 20h (B) or 44h (C). FDR-q values were generated using a 1000-geneset permutations. Log_2_ TPM expression magnitude of indicated genes is shown as a mean, 75% confidence interval (box) and 95% confidence interval (whisker) as well as individual data points, comparing FO B cells on OP9 + LPS (black) to OP9-DL1 + LPS (green) (D). RNA sequencing reflects one experiment with 3 technical replicates for each condition.

### MZ B cells are uniquely responsive to TLR3

One earlier report suggests that among all peripheral B cells only MZ B cells express mRNA encoding TLR3^32^. To explore the possibility that MZ B cells can respond to TLR3 ligation, we stimulated purified follicular or MZ B cells with graded doses of the TLR3 agonist Poly (I:C)^39^. A large fraction of MZ B cells underwent one round of cell division (**Figure 5A**) and plasma cell differentiation in response to Poly (I:C) stimulation, even to low doses (**Figure 5B**), whereas we detected no proliferation or differentiation of Poly(I:C)-stimulated follicular B cells. Indeed, MZ B cells underwent division and yielded both IgM and IgG-producing plasma cells above background with as little as 0.001 μg/ml Poly (I:C). Also, we questioned whether Notch2-independent B1 B cells would respond to TLR3 signals given they share many functional characteristics with MZ B cells, such as their innate-like poising and hyperresponsiveness to TLR signals. Indeed, we failed to observe evidence for TLR3-responsiveness for peritoneal cavity B1 B cells (**Extended figure 4A**). To evaluate whether *Tlr3* expression in MZ B cells is Notch2-dependent, we evaluated our published RNAseq data generated with purified follicular and MZ B cells harvested shortly after inoculating adult B6 mice with a Notch2 blocking antibody^12,40^. As shown in **Figure 5C**, for MZ B cells *Tlr3* transcripts decreased after only 12 hours of Notch2 blockade, whereas for follicular B cells *Tlr3* transcripts were substantially lower and unaffected by Notch2 blockade. Furthermore, *Myd88* transcript abundance was modestly decreased after 48 hours, whereas mRNA levels for *Ticam1*, the gene encoding TRIF, trended modestly higher in MZ B cells but was not impacted by Notch2 blockade. Transcript abundance for *Cd180*, encoding the LPS coreceptor RP105, was higher in MZ B cells than in follicular B cells and reduced rapidly upon blockade of Notch2 signaling. Together, these data highlight potential mechanisms through which Notch2 can regulate TLR responses. Including for TLR3

**Figure 5.**
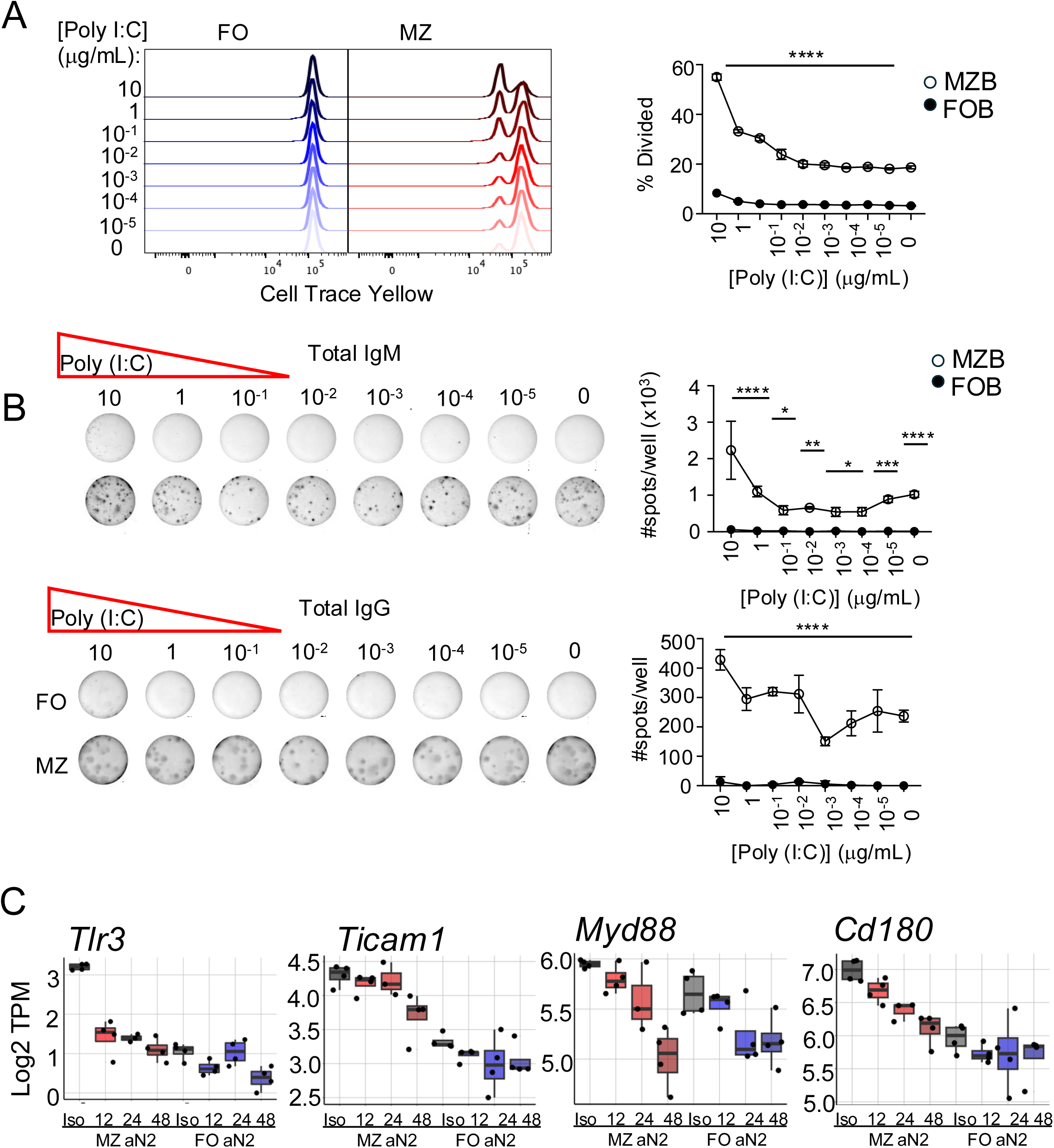
TLR3-responsive program in resting MZ B cells is established by Notch2. Purified Follicular or MZ B cells were examined for receptiveness to TLR3 agonist Poly (I:C), depicted by Cell Trace Yellow (CTY) dilution for follicular (left) or MZ (right) B cells. Poly (I:C) dose depicted to the left of the plots in μg/mL. Results are quantified in the graph to the right (A). ELISpots from titrating Poly (I:C) in follicular or MZ B cell cultures were performed 48 hrs after stimulation, and total IgG+ spots/well were measured and quantified (right), the 1:24 dilution is shown (left). Mice were treated with monoclonal blocking antibody against Notch2 (aN2) or isotype control (iso) for 12-48 hours, and MZ (red) or FO (blue) cells were subsequently sorted from the spleens of those mice for bulk RNA sequencing. Transcript levels for *Tlr3*, *Ticam1*, *Myd88*, and *Cd180* are shown on a Log2 transcripts per million (TPM) scale, displayed as a mean with a 75% confidence interval (box) and 95% confidence interval (whisker) (C). 2-way ANOVA performed A-B, **** = p<0.0001. Triplicate cultures, repeated in at least 5 independent experiments. RNA sequencing reflects one experiment with three biological replicates.

### Notch2 regulates TLR3 responsiveness

To examine whether Notch2 signals alone were sufficient to instruct a TLR3 response program, we cultured follicular B cells on OP9 or OP9-DL1 with varying amounts of Poly (I:C). Follicular B cells did not respond to Poly (I:C) in coculture with OP9 control cells. In contrast, DL1-stimulated follicular B cells experienced extensive cell division, beginning as early as 42 hours post-stimulation with upwards of 7 divisions by 90 hours with 20μg/ml Poly (I:C) (**Figure 6A**). This result was supported by the increased total cell numbers recovered from culture over time, both among live (left) and total (right) cells, particularly at 66h and beyond (**Figure 6B**). We additionally observed a decrease in the percentage of surviving cells over time, likely relating to the increase in cell expansion for Notch2-experienced cells (**Figure 6C)**. Furthermore, kinetic analysis of MDNs among responders (left) and all B cells (right) in OP9-DL1 cultures suggested that LPS and Poly (I:C)-stimulated cells divided with similar timing and magnitude (**Figures 3B, 6D**). Linear regression analysis of the MDN revealed a substantially faster time to first division and subsequent cycle time (62.8h vs 507h and 36.6h v 478h, respectively) signifying that not only does Notch2 shorten the time to divide for B cells as it does for LPS, but it permits B cells to respond to TLR3 signals altogether (**Figure 6E**). As expected, DL1-facilitated responsiveness to Poly (I:C) was abrogated by inhibition of the key adaptor TRIF using Pepinh-TRIF TFA, a 30-amino-acid peptide that blocks the TRIF-TLR interaction directly^41^ (**Figure 6F**). On the other hand, for other endosomal nucleic acid sensors TLR7 and TLR9, DL1 appeared to have a minimal effect on the overall proliferative capacity over time, Myc induction, and time to divide that was significantly different only at early time points (**extended Figures 1 & 2**). Thus, we conclude that transient DL1 exposure is sufficient to turn on a TLR3 program and render B cells responsive to this signal in a TRIF-dependent manner.

**Figure 6.**
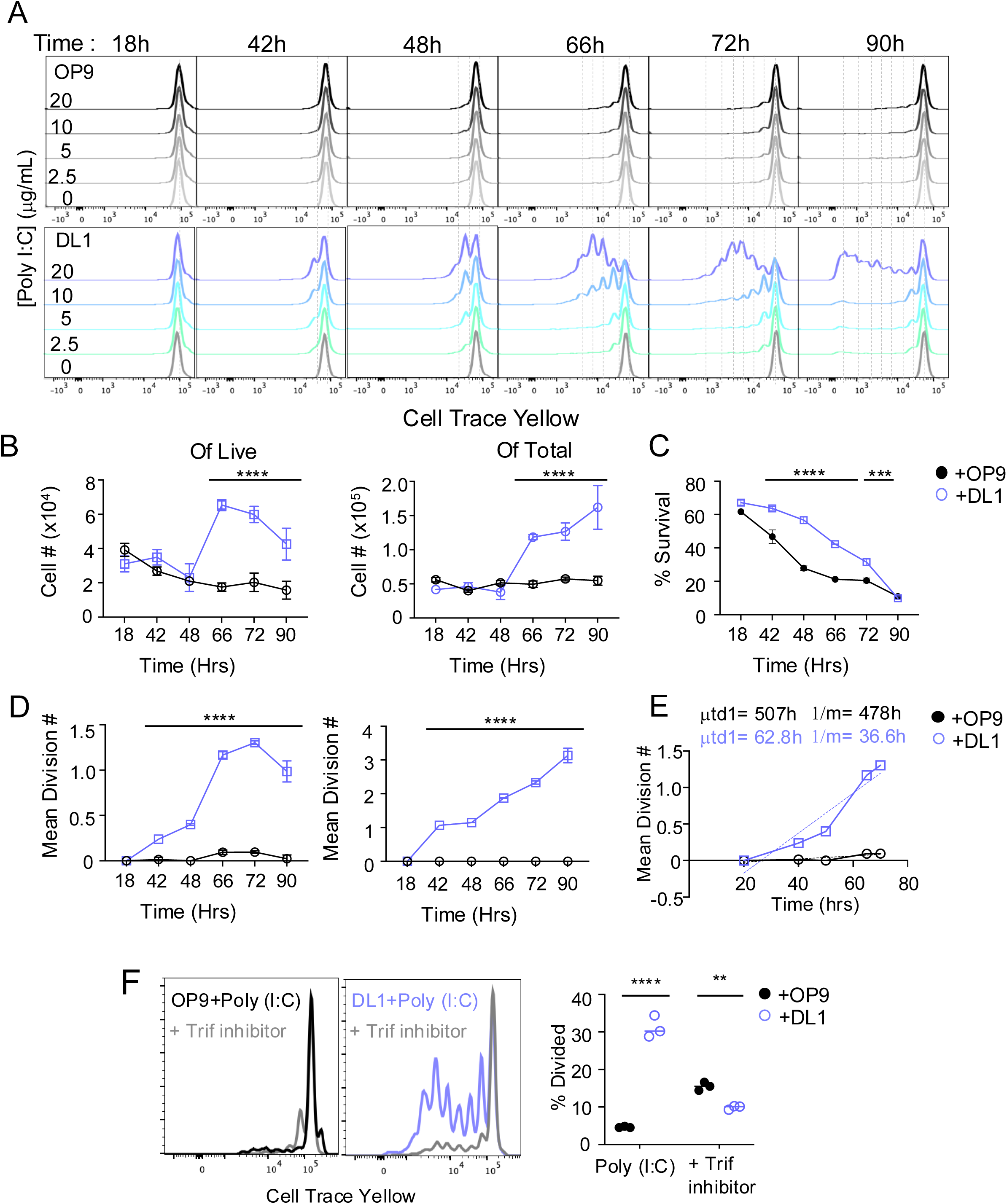
Forced Notch2 signals instruct a TRIF-dependent TLR3-responsive program for FO B cells. Purified FO B cells were Cell Trace Yellow labeled and co-cultured with OP9 stromal cells (grey) or OP9 stromal cells transduced to express Notch2-ligand DL1 (teal) and stimulated with indicated amounts of Poly (I:C) in culture for 18-90 hours with the concentration denoted by increasingly darker shades of teal or grey. The resulting histograms are depicted (A). All graphs display stimulation with the 20 μg/mL dose. Cell number over time from live (left) and total (right) from cultures shown (B). % of ToPro-3^-^ cells over time in culture depicted between OP9 (black) and OP9-DL1 (violet) (C). Resulting mean division numbers (MDN) over time are depicted for total cells (left) or responding cells (right). (D). Division rate is depicted, with simple linear regression analysis performed on the linear portion of the MDN plot for the highest dose between OP9 and DL1 conditions, time to first division was found to be 507h and 62.8h and time to complete subsequent divisions was found to be 478h and 36.6h for OP9 and DL1, respectively (E). Poly (I:C) proliferative response is depicted through Cell Trace Yellow histograms for OP9 (left) and DL1 (right) +/-40 μg/mL of Trif inhibitor, percent divided depicted to the right (F). 2-way ANOVA performed B-D,F * = p<0.05, ** = p<0.005, *** = p<0.001 **** = p<0.0001. Triplicate cultures, repeated in at least 5 independent experiments.

### A role for BTK in Notch2-fostered B cell responses to Poly (I:C)

To elucidate signaling pathways used by DL1-stimulated B cells to respond to TLR3 signals, we sought to extend past work showing that TLR3 signaling in macrophages requires the Tec kinase BTK^28^. Resting MZ B cells possessed higher levels of phosphorylated BTK compared to follicular B cells (**Figure 7A**). Similarly, unstimulated FO B cells on DL1 after 72 hours exhibited higher levels of phosphorylated BTK, suggesting that Notch2 signaling augments phosphorylation of BTK (**Figure 7B**). We therefore compared the impact of compounds that inhibit BTK (Ibrutinib) versus other canonical mediators of proximal B cell receptor signaling such as Lyn (I406) and Syk (R406, R788) on Poly (I:C)-and LPS-driven follicular B cell responses in OP9 and OP9-DL1 cultures. All inhibitors were first titrated to identify the lowest point with which B cell proliferation in response to receptor cross-linking was inhibited with minimal cell death to identify the optimal concentration of each compound (**Extended Figure 3B, 3C**). Ibrutinib (Btk *i*) profoundly inhibited DL1-dependent Poly (I:C)-driven B cell proliferation without affecting LPS-stimulated proliferation, whereas Syk and Lyn inhibitors had no effect on Poly (I:C)-or LPS-driven proliferative responses (**Figure 7C**). Likewise, Ibrutinib inhibited Poly (I:C)-driven but not LPS-driven plasma cell differentiation (**Figure 7D**). We further validated that Poly (I:C)-driven proliferation was due to TLR3 activity alone by crossing TLR3^f/f^ mice to an Mb1cre background, thereby preventing TLR3 expression in all B cells. We observed a complete loss of Notch2-driven proliferation in response to Poly (I:C) in TLR3^f/f^; Mb1cre het mice mice compared to the robust proliferative response observed in the Mb1cre het control (**Figure 7E**). We conclude that Notch2-augmented phosphorylated BTK permits TLR3 activation in B cells. We then examined B cells from CBA.xid mice which possess an X-linked null mutation in the BTK gene^42^. BTK-null MZ B cells exhibited a substantial loss of both proliferation and plasma cell differentiation in response to Poly (I:C) but not LPS (**Extended Figure 4 B, 4C**).

**Figure 7.**
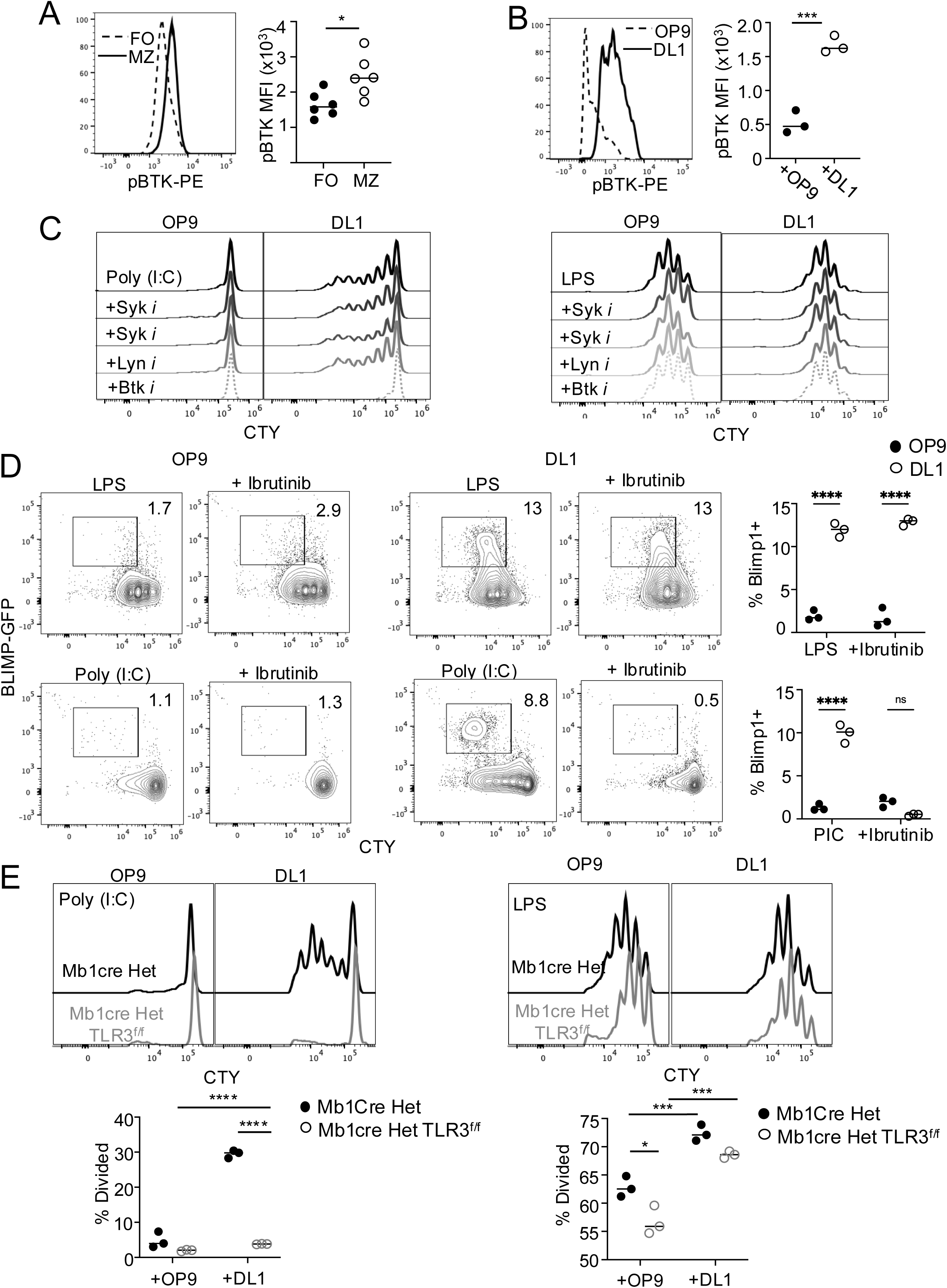
Phosphorylation of BTK is augmented by Notch2 and required for Poly (I:C) responsiveness. Phosphorylated Btk was examined by flow cytometry comparing follicular (dashed) and MZ B cells (solid), and quantified to the right (A). Flow cytometry histograms display phosphorylated Btk among resting follicular B cells after 72h on OP9 (dashed) or OP9-DL1 (solid) and are quantified to the right (B). Purified FO B cells in the OP9 or OP9-DL1 culture system were tested against various B cell receptor signaling inhibitors in the presence of TLR3 (left) or TLR4 (right) activation. Cell trace histograms reveal B cells stimulated with 20 μg/mL Poly (I:C) +/-inhibitors to Syk (R788 and R406), Lyn (I406), or Btk (Ibrutinib) that were previously tested and titrated to determine the optimal dose for each (C). Plasma cell induction from BLIMP1-GFP reporters at 72 hours for LPS +/-Ibrutinib (top) or Poly (I:C) +/-Ibrutinib (bottom) stimulated cells on OP9 or DL1is displayed, numbers depict % of Blimp1+ cells (D). Histogram plots depict cell trace histograms of Poly (I:C) (left) or LPS (right)-stimulated follicular B cells from Mb1cre Het (black line) or Mb1cre het TLR3^f/f^ mice (grey line) on OP9 (left) or OP9-DL1 (right), percent divided quantified to the right. * = p<0.05, *** = p<0.001 ****, = p<0.0001 based on 2-way ANOVA. Results were repeated in 3-4 independent experiments.

### Notch2 drives plasma cell differentiation and class-switching independent of T cell help

Given the plasma cell-poised nature of MZ B cells, we next asked whether DL1 was sufficient to foster plasma cell differentiation in response to TLR3 stimulation. Poly (I:C)-stimulated follicular B cells generated substantial numbers of Blimp1-GFP+ plasma cells over time in concert with DL1 in a TRIF-dependent manner (**Figure 8A, extended Figure 3**). Because our past work showed that DL1-stimulation augmented class switching due to CD40 ligation^43^, we also performed ELISpot assays to quantify cells secreting each IgG subclass after co-stimulation with DL1 and either LPS or Poly (I:C). As shown in **Figure 8B**, we observed substantial numbers of total IgG, IgG_2b_ and IgG_2c_-secreting plasma cells induced by Notch2 activity in response to both LPS or Poly (I:C). Of note, we examined activation of other TLRs expressed in B cells and neither R848 (TLR7) or CpG (TLR9) stimulation induced plasma cell differentiation with or without DL1 (**Extended figures 1F,2F**). These results indicate that DL1-augmented TLR3 and TLR4 responses are uniquely sufficient to drive IgG class-switched plasma cell differentiation independent of canonical T cell-derived signals, particularly for IgG_2b_ and IgG_2c_.

**Figure 8.**
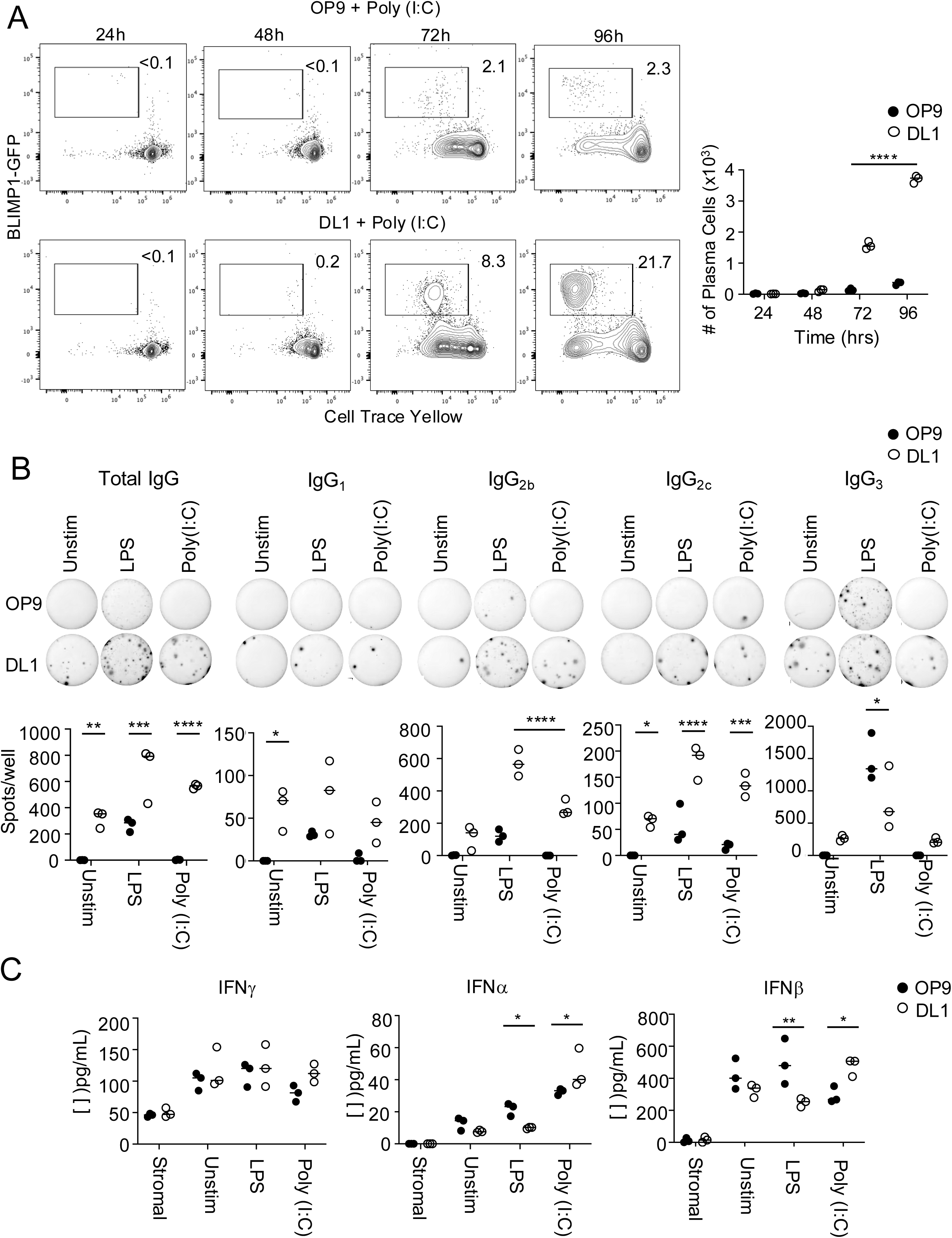
Forced Notch2 augments plasma cell differentiation and IgG2b,2c class switching. Purified FO B cells from BLIMP1 ^+/GFP^ mouse splenocytes were Cell Trace Yellow labeled and co-cultured with OP9 or DL1. Over time in culture in addition to 10 μg/mL Poly (I:C) plasma cell differentiation was tracked by the BLIMP1-GFP+ population, numbers depict percent of positive cells. Quantitated are the absolute number of BLIMP1+ cells over time on OP9 (solid black circles) or DL1 (open black circles) (A). ELISpots were performed to detect the number of antibody secreting cells in culture with OP9 (top) or DL1 (bottom) +/-LPS or Poly (I:C) as indicated. Total IgG, IgG_1_, IgG_2b_, IgG_2c_, and IgG_3_ were measured after 48h in culture with indicated stimuli. 1:24^th^ dilution from the total cells in the well are displayed, average spots per well quantified below (B). IFNs were measured by flow cytometric cytokine bead arrays from tissue culture supernatants and quantified. FO B cells were co-cultured on OP9 or OP9-DL1 for 72 hours alone (unstim) or stimulated with LPS or Poly (I:C), with stromal cells alone being tested for secretion as well (stromal). Cytokine concentrations were quantified using the BioLegend cytokine bead array analysis platform (C). 2-way ANOVA performed * = p<0.05, ** = p<0.005, *** = p<0.001 **** = p<0.0001. Triplicate cultures, repeated in at least 3 independent experiments.

Of note, we did not observe increased levels of IFNγ with OP9-DL1 coculture. Alternatively, we observed a very modest increase in type I interferons α and β on DL1 following Poly (I:C), whereas with LPS stimulation they decreased compared to OP9 controls (**Figure 8C**). Together these data uncover a unique plasma cell and class-switching program turned on by Notch2 signals in the presence of TLR3 and TLR4 stimulation independent of cognate help and IFNγ.

## Discussion

Much of what we know about Notch signaling within the immune system pertains directly to development and guiding cell fate decisions. For B cells, Notch2 is the signal that guides MZ B cell fate commitment. Fewer studies however have investigated the consequences of Notch2 signaling beyond development. Here, we describe a novel role for Notch2 to endow frontline B cells with unique functional characteristics such as more rapid division, increased proliferative activity, PC differentiation potential, class-switching independent of T cell help, and cytokine secretion – all relevant to the rapid control of blood-borne bacterial and viral infections.

We demonstrated that naïve follicular B cells can be endowed with a hyperresponsiveness to LPS upon transient Notch2 signaling, leading to both augmented division and PC differentiation. Notch2-dependent MZ B cells have long been known for their unique hyperresponsiveness to blood-borne encapsulated bacteria, with emphasis on both TLR4 and poly-reactive antigen receptors^10,8,11^. These hallmarks are complemented by the specialized localization of MZ B cells; the spleen filters upwards of 5% of the cardiac output, a large and continuous blood supply with a low flow rate as it transits through the red pulp and the marginal sinus. This design serves to maximize antigen interactions and enable constant surveillance of blood-borne antigens in the marginal zone^44,45^. MZ B cells being endowed with hypersensitivity to TLR signals is consistent with their frontline positioning and specialized roles. This space is additionally occupied by a complex network of specialized macrophages, dendritic cells, and neighboring lymphoid follicles harboring Notch-ligand expressing fibroblastic reticular cells^16,44^. This immune network affords access to DL1, which is imperative for both the survival and maintenance of MZ B cells^18,12^. We demonstrate here that continuous Notch2 signaling additionally establishes poising for TLR4 responses.

Most notably, we found that access to DL1 turned on a TLR3 response program triggered by double stranded RNA. It has been reported that MZ B cells uniquely among B cells express transcripts for *Tlr3*, however it has never been determined what establishes this unique signaling axis. Here, we tie its functionality in B cells to Notch2 signaling. Within peripheral lymphoid tissues, red pulp macrophages, CD8+ dendritic cells, and MZ B cells are among the highest expressors of TLR3, all localized within this distinct anatomical niche^46^. Notably, MZ B cells and subsets of CD8+ dendritic cells are reliant on Notch2 signaling for their development^47^. While it is unclear whether red pulp macrophages rely on Notch signaling as well, they express Notch2 as highly as MZ B cells as well as several Notch2 targets such as Hes1^46^. It is widely accepted that many common viral infections producing systemic symptoms (i.e. fever, chills, nausea, etc.) result in viremia, or blood-borne systemic viral infection^48^. It is therefore conceivable that MZ B cells would play a role in the rapid ‘innate-like’ detection and response to blood-borne viremia in this critical niche just as they would for bacterial antigens. Indeed, several studies highlight the need for both ‘innate-like’ B-1 B cells and extrafollicular responses early on in viral infections such as influenza^49^. However, there exists no study on the direct relationship between MZ B cells and viral antigens through TLR3 signaling. In addition to MZ B cells’ responsiveness, we demonstrated that Notch2 signals can endow previously unresponsive follicular B cells with a TLR3 response program, as co-culture on OP9-DL1 led to robust proliferation to the TLR3 agonist Poly (I:C) after only 42-48 hours. To our knowledge, this is the first report of a link between rapid TLR-responsiveness and Notch2 experience, emphasizing a potential role for MZ B cells in innate viral sensing.

We demonstrated that transient Notch2 experience changes the landscape of phosphorylated signaling molecules that may alter a variety of B cell response programs. Resting MZ B cells exhibited higher baseline levels of phosphorylated BTK compared to follicular B cells. After transient culture on Notch2 ligands however, phosphorylation of BTK increased significantly in follicular B cells. BTK, a signaling adapter typically involved in antigen receptor signaling, has been reported to aid in phosphorylation TLR3 itself within endosomes^28^. Here we demonstrate the Notch2-instructed Poly (I:C) response in follicular B cells is dependent on BTK, as Ibrutinib shuts down the ability of folli cular B cells to proliferate or differentiate.

These results establish that Notch2 is involved in augmenting BTK signaling, with an unforeseen consequence of targeting TLR3. Further work however is needed to gain understanding of how this axis is established and whether B cells are required for TLR3 responses. Indeed, reports have demonstrated crosstalk between antigen receptor components and TLR adapters in signalosome complexes^50^. These findings have profound implications for targeting and augmenting B cell responses in vaccine settings using TLR3 adjuvants, for one.

These results establish that Notch2 is involved in augmenting BTK signaling, with an unforeseen consequence of targeting TLR3. Further work however is needed to gain understanding of how this axis is established and whether B cells are required for TLR3 responses. Indeed, reports have demonstrated crosstalk between antigen receptor components and TLR adapters in signalosome complexes^50^. These findings have profound implications for targeting and augmenting B cell responses in vaccine settings using TLR3 adjuvants.

We elucidated several distinct consequences of forced Notch2 signaling on B cells; among them are the robust amplification of cell division by both timing and magnitude. We employed mathematical models adapted from the Hodgkin group^37^ to reveal that sustained Notch2 can augment proliferative responses to TLR signals by shortening the time to first division, accelerating entry into G1, and increasing both the resulting cell number and mean division number of responding B cells. These findings were accompanied by an earlier induction of the mitotic regulator Myc. These results are profound, as prior studies have implicated the time to first division for lymphocytes is ‘fixed’ among responding cells. Further, this process is said to be largely stochastic, i.e. not governed by any cell-extrinsic events^33,51^. Here, we demonstrated that an otherwise fixed process for dividing lymphocytes can be influenced by one cell-extrinsic factor, but how this reprograming is achieved is still elusive. Several possible outcomes of sustained Notch2 signals are targeting of transcriptional regulators for cell division, in addition to Myc, such as cyclin-dependent kinases and mTORC1. Accelerating the division clock for B lymphocytes is a impactful consequence that could stand to enhance effector responses to a variety of signals. Though Notch2 has been implicated in altering cell division before, for example in the case of cardiomyocytes^52^, this is the first study to highlight its role in driving and accelerating mitotic programs for B cells.

Finally, we observed that Notch2 signals endow B cells with a plasma cell differentiation potential independent of T cell help. More recent evidence points to MZ B cells’ robust transcriptional poising for plasma cell differentiation^12^. These findings connect sustained Notch2 signaling to differentiation potential and have elucidated a mechanism for the rapid thymus-independent plasma cell program MZ B cells are known to exhibit. Transient Notch2 experience alongside a TLR ligand can unleash a plasma cell program independent of cognate help in follicular B cells. Building off our prior findings in the lab, Notch2 may achieve this through plasma cell-related gene programs (Xbp1, Blimp1, UPR, etc.) as well as targeting of mTORC1 to aid in cell growth and expansion of endoplasmic reticulum to gear up for mass protein production^53^. The following were validated in our RNA sequencing results, showing an early transcriptional signature for plasma cell poising in the presence of Notch2 signals.

Though these rapid T cell-independent plasma cells are classically thought of as making primarily IgM, we also noted a significant portion of IgG-positive plasma cells, particularly those making IgG_2b_ and IgG_2c_. These findings validate earlier observations that Notch2 can augment CSR^43^. Previous reports characterizing CD40 deficiency in mice also demonstrate an intact ability for B cells to class-switch to IgG in a thymus-independent fashion, validating the idea that IgG-class switching can indeed occur in absence of cognate help for bacterial antigens^54^.

Together these findings underscore the plasticity and heterogeneity among mature B cells, which are otherwise often classified as distinct in their phenotype and function. Whereas MZ B cells are classically characterized as a uniquely ‘innate-like’ subset of mature B cells, ultimately many of these features of poised cell states can be ascribed to any B cell in the context of transient Notch2-experience. These results underscore how a complex immune system functions to control an immune response in layers, with highly conserved fail-safe mechanisms to ensure that an innate-like pool of cells is ready to fight blood-borne infections effectively and rapidly.

## Acknowledgments

The authors thank Drs. Michael Cancro, Warren Pear, and Avinash Bhandoola for helpful comments and profound wisdom. This work was supported by National Institutes of Health R21-AI174069 and R01-AI175185 (D.A.), R01-AI091627 and R01-CA278976 (I.M.) and 1F31 AI179166 (J.L.), T32 AI007632 (JL), and T32 CA009171 (N.J.T.). D.B.W. is supported by NIH P01AI165066, NIH/NIAID Collaborative Influenza Vaccine Innovation Centers contract 75N93019C00051, and INOVIO Pharmaceuticals SRA 21-05. Additional funding to D.B.W provided by the W.W. Smith Charitable Trust Distinguished Professorship in Cancer Research and The Jill and Mark Fishman Foundation.

## Author contributions

J.L. designed and performed experiments, analyzed data and wrote the manuscript, I.R. and B.T.G. analyzed data and provided figures, N.J.T. evaluated assays, D.W. and I.M. reviewed data and supervised the work, D.A, selected the research questions, supervised the work, and wrote the manuscript.

