## Supplemental Information for "Notch2 signaling instructs viral and bacterial TLR responsiveness in B cells"

Fig S1

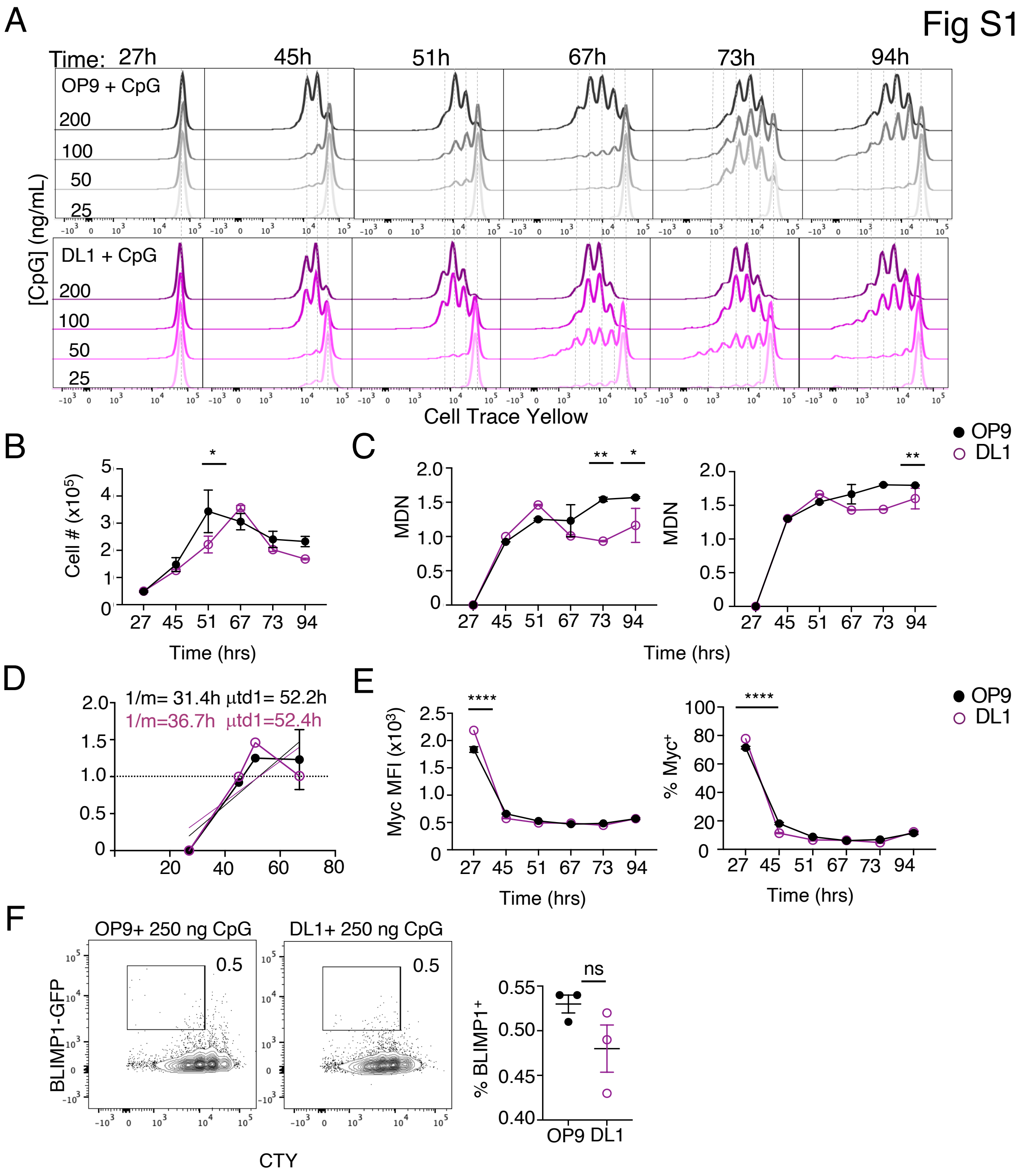
A
Time: 27h 45h 51h 67h 73h 94h
OP9 + CpG
DL1 + CpG
200
100
50
25
[CpG] (ng/mL)
Cell Trace Yellow
B
Cell # (x10^5)
Time (hrs)
C
MDN
Time (hrs)
OP9
DL1
D
1/m= 31.4h µtd1= 52.2h
1/m=36.7h µtd1=52.4h
E
Myc MFI (x10^3)
% Myc+
Time (hrs)
F
OP9+ 250 ng CpG
DL1+ 250 ng CpG
BLIMP1-GFP
CTY
% BLIMP1+
ns
OP9 DL1

**Supplemental Figure 1**. **Forced Notch2 signals have minimal impact on TLR9 response.** FO B cells were purified from cMyc-GFP reporters and Cell Trace Yellow labeled and co-cultured with OP9 stromal cells (grey) or OP9 stromal cells transduced to express Notch2-ligand DL1 (magenta) and stimulated with indicated amounts of CpG in culture for 27-94 hours, concentration is denoted by increasingly darker shades of magenta or grey. Resulting histograms are depicted (A). All graphs display stimulation with the 200 ng/mL dose. Total cell number over time from cultures is shown (B). Resulting mean division numbers (MDN) over time are depicted for total cells (left) or responding cells (right) (C). Division rate is depicted, with simple linear regression analysis performed on the linear portion of the MDN plot for the highest dose between OP9 and DL1 conditions, time to first division was found to be 52.2h and 52.4h and time to complete subsequent divisions was found to be 31.4h and 36.7h for OP9 and DL1, respectively (D). Resulting cMyc-GFP MFI and % Myc+ cells over time are quantified (E). BLIMP1+ plasma cells are quantified in culture vs Cell Trace Yellow after 72h in culture with 250 ng CPG, frequency of BLIMP1+ quantified to the right (F). 2-way ANOVA performed B-D,F * = $p<0.05$, ** = $p<0.005$, **** = $p<0.0001$. Triplicate cultures, repeated in at least 5 independent experiments.

Fig S2

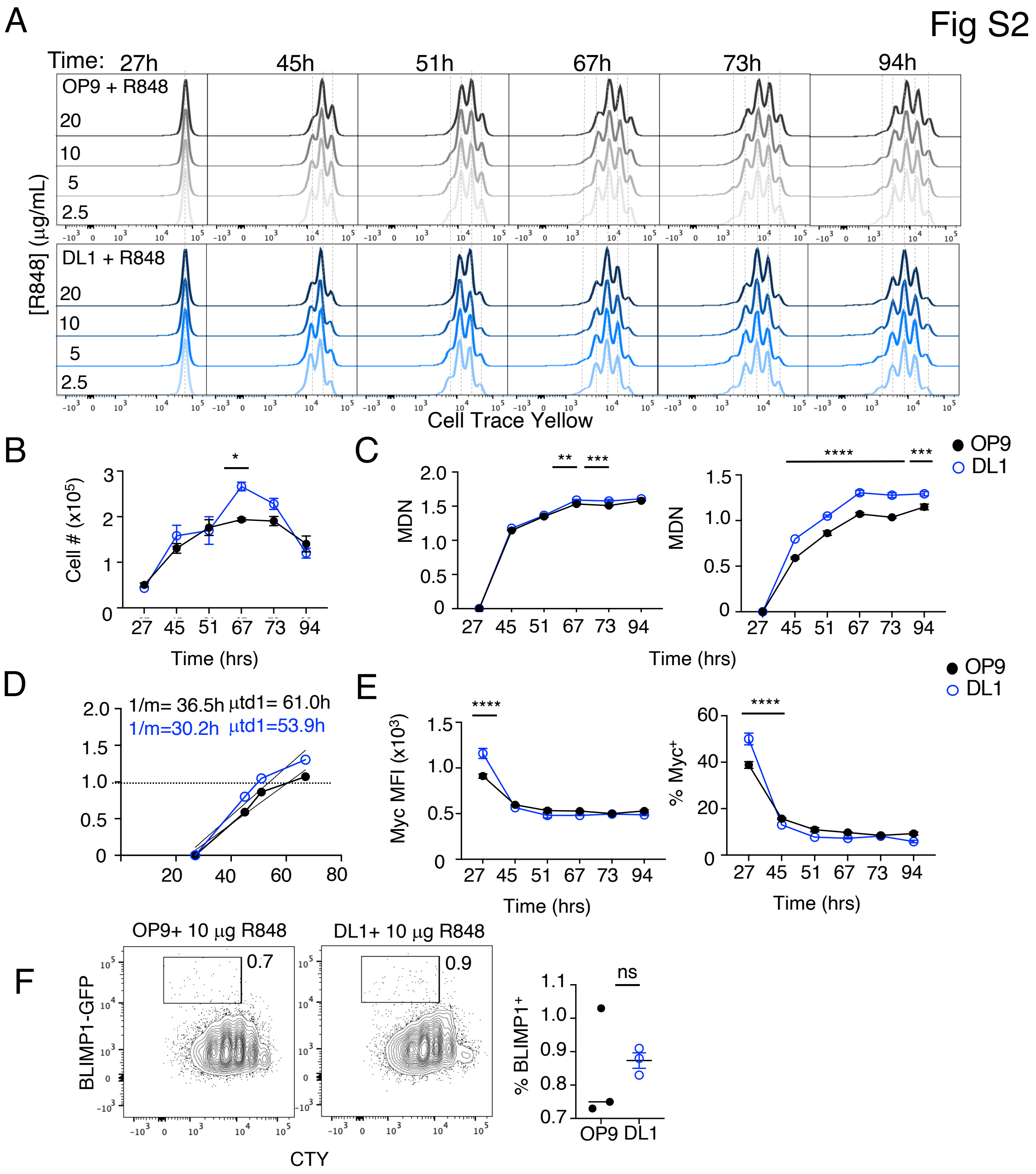

A
Time: 27h 45h 51h 67h 73h 94h
OP9 + R848
DL1 + R848
20
10
5
2.5
[R848] (μg/mL)
Cell Trace Yellow
B
Cell # (x10⁵)
Time (hrs)
C
MDN
Time (hrs)
OP9
DL1
D
1/m= 36.5h μtd1= 61.0h
1/m=30.2h μtd1=53.9h
E
Myc MFI (x10³)
% Myc⁺
Time (hrs)
F
OP9+ 10 μg R848
DL1+ 10 μg R848
0.7
0.9
BLIMP1-GFP
CTY
% BLIMP1⁺
ns
OP9 DL1

**Supplemental Figure 2**. **Forced Notch2 signals have minimal impact on TLR7 response.** FO B cells were purified from cMyc-GFP reporters and Cell Trace Yellow labeled and co-cultured with OP9 stromal cells (grey) or OP9 stromal cells transduced to express Notch2-ligand DL1 (blue) and stimulated with indicated amounts of R848 in culture for 27-94 hours, the concentration denoted by increasingly darker shades of blue or grey. Resulting histograms are depicted (A). All graphs display stimulation with the 20 μg/mL dose. Total cell number over time from cultures shown (B). Resulting mean division numbers (MDN) over time are depicted for total cells (left) or responding cells (right) (C). Division rate depicted, with simple linear regression analysis performed on the linear portion of the MDN plot for the highest dose between OP9 and DL1 conditions, time to first division was found to be 61h and 53.9h and time to complete subsequent divisions was found to be 36.5h and 30.2h for OP9 and DL1, respectively (D). Resulting cMyc-GFP MFI and % Myc+ cells over time are quantified (E). BLIMP1+ plasma cells quantified in culture vs Cell Trace Yellow after 72h in culture with 10 μg R848, frequency of BLIMP1+ quantified to the right (F). 2-way ANOVA performed B-D,F * = $p<0.05$, ** = $p<0.005$, *** = $p<0.001$ **** = $p<0.0001$. Triplicate cultures, repeated in at least 3 independent experiments.

Fig S3

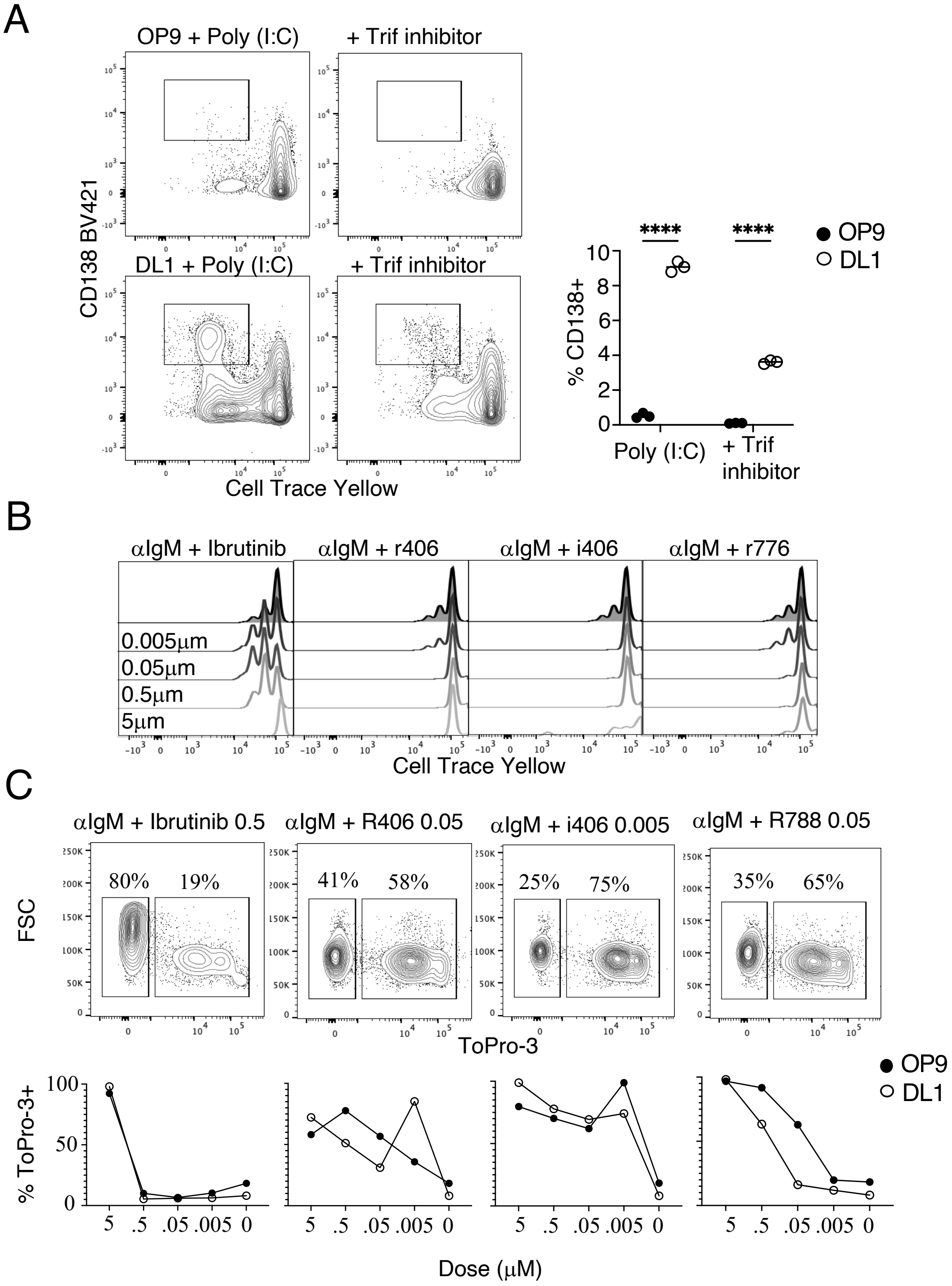
A
OP9 + Poly (I:C)
+ Trif inhibitor
DL1 + Poly (I:C)
+ Trif inhibitor
CD138 BV421
Cell Trace Yellow
****
****
OP9
DL1
% CD138+
Poly (I:C)
+ Trif inhibitor
B
αIgM + Ibrutinib
αIgM + r406
αIgM + i406
αIgM + r776
0.005μm
0.05μm
0.5μm
5μm
Cell Trace Yellow
C
αIgM + Ibrutinib 0.5
αIgM + R406 0.05
αIgM + i406 0.005
αIgM + R788 0.05
80%
19%
41%
58%
25%
75%
35%
65%
FSC
ToPro-3
OP9
DL1
% ToPro-3+
Dose (μM)

**Supplemental Figure 3. Various inhibitors tested against Poly (I:C) and F(ab’2) anti-IgM.** Purified FO B cells were co-cultured with OP9 or DL1 and stimulated with 10 μg/mL Poly (I:C) for 72 hrs in the presence or absence of 40 μg/mL Trif inhibitor. Plasma cell differentiation is examined by CD138 staining, percent positive is quantified to the right (A). Inhibitors to Btk, Syk, and Lyn were tested and titrated against BCR ligation with 10 mg/mL F(ab’2) anti-IgM to determine the optimal dose that inhibits proliferation (B) with minimal cell death measured by ToPRO-3 labeling (C). ToPRO-3 plots quantified at each dose of inhibitor below. 2-way ANOVA performed (A), **** = $p<0.0001$.

### Fig S4

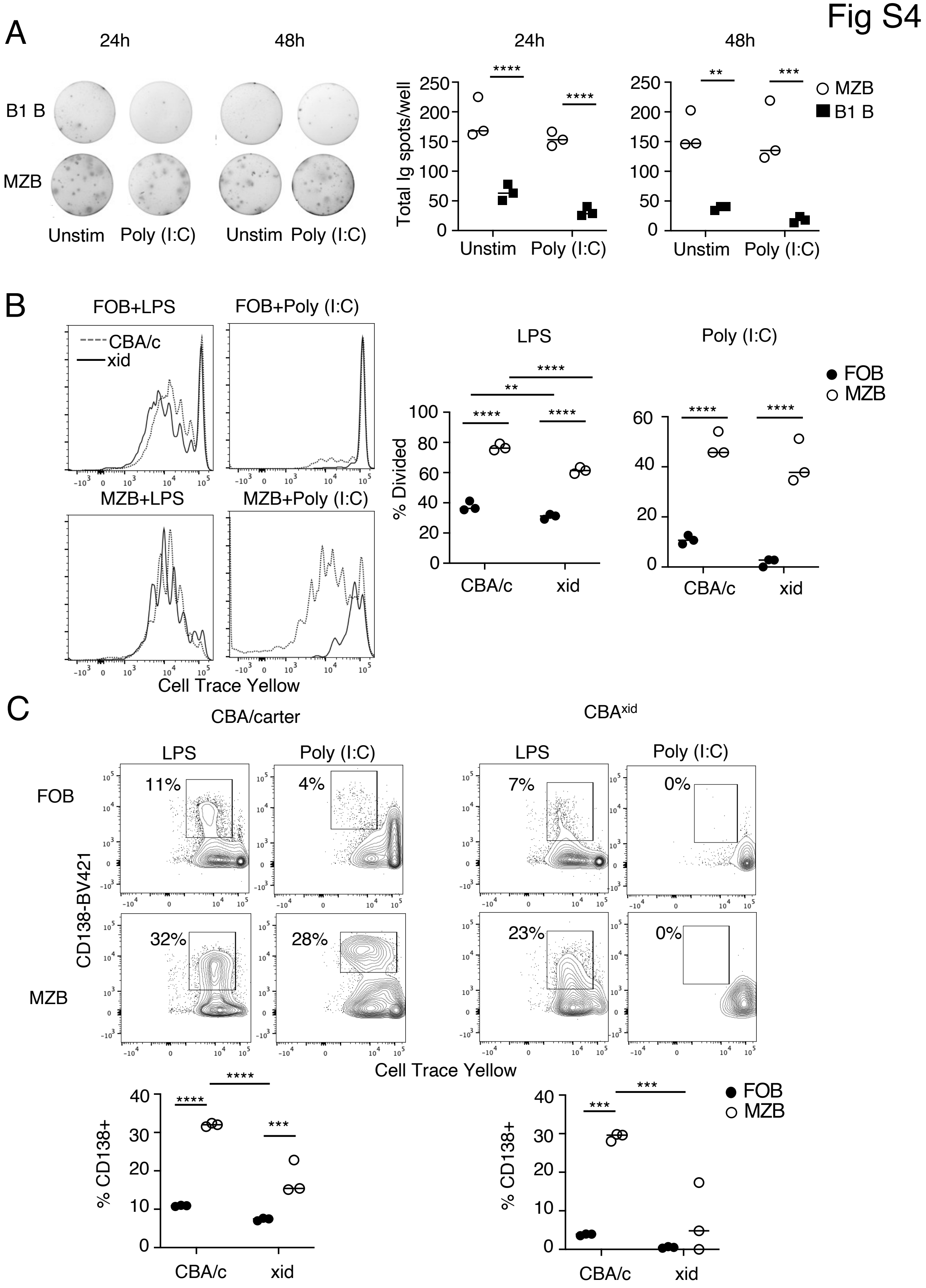
A
24h
48h
B1 B
MZB
Unstim
Poly (I:C)
Total Ig spots/well
**
***
****
MZB
B1 B
B
FOB+LPS
FOB+Poly (I:C)
MZB+LPS
MZB+Poly (I:C)
CBA/c
xid
Cell Trace Yellow
LPS
Poly (I:C)
% Divided
FOB
MZB
C
CBA/carter
CBAxid
CD138-BV421
11%
4%
7%
0%
32%
28%
23%
0%
% CD138+

**Supplemental Figure 4. MZB cell TLR3 and 4 responses rely on BTK, other mediators appear unaffected by DL1 experience.**

B1 B or MZB cells were cultured for 24 or 48h alone or with Poly (I:C) and ELISPOTs were performed to measure total Ig spots per well (of the tissue culture plate) and are quantified to the right (A). Follicular or Marginal zone B cells were purified from the spleens of CBA/carter mice or $CBA^{xid}$ mice and stimulated with LPS or Poly (I:C). Cell trace histograms depict division after 72 hours with CBA/carter controls (dashed line) or $CBA^{xid}$ (solid line). % of resulting division is quantified to the right (B). Plasma cell differentiation was measured by CD138 staining for CBA/carter (left) or $CBA^{xid}$ mice (right) stimulated with LPS or Poly (I:C), % positive quantified below (C). 2-way ANOVA performed (A-C), ** = p<0.005, *** = p<0.001, **** = p<0.0001. Results repeated in at least 3 independent experiments.
